# Rising daptomycin resistance in *Enterococcus faecium* across a hospital system occurred via rampant recurrent evolution and occasional transmission between patients

**DOI:** 10.1101/2023.05.09.540070

**Authors:** Robert J. Woods, Meghan Forstchen, Clare Kinnear, Jordan McKaig, Twisha Patel, Kevin Tracy, Carol Young, Andrew F. Read

## Abstract

The rise of antibiotic resistance in a population involves two distinct processes: the origin of resistance and its spread. Here we study the contribution of both processes to the increase in daptomycin resistance in *Enterococcus faecium* in a hospital system. This case-control genomic study includes whole-genome sequencing of 82 isolates obtained from 24 case patients with clinically determined daptomycin-resistance and 24 controls. Among the case patients, the first isolate was resistant in 15 patients (R patients) while in the remaining nine the first isolate was susceptible but was followed by one or more resistant isolates (SR patients). Mutations in a set of candidate daptomycin resistance genes were compared within and between all patients. Additionally, among closely related isolates, mutations were identified across the entire assembled genome. Daptomycin resistance evolved separately multiple times and there was no phylogenetic clustering of the R or the SR groups. Six of the nine SR pairs gained mutations in previously identified candidate loci for daptomycin resistance, with the major cardiolipin synthase (*clsA*) being mutated most frequently. The hospital-wide increases in daptomycin resistance in *E. faecium* was the result of recurrent evolution taking multiple evolutionary pathways and occasional transmission of resistant isolates between patients.

**Importance:** Antimicrobial resistance in healthcare settings presents an important challenge, because infections with resistant organisms are associated with higher cost, longer hospital stays and worse outcomes for patients. However, it can be difficult to identify the factors driving the increase in resistance, specifically the relative contribution of resistance arising anew through mutation versus the transmission of resistant organisms from patient to patient. We study a hospital where resistance to daptomycin was increasing among *Enterococcus faecium*, an important hospital pathogen. We find the increase in resistance was the results of resistance arising many times independently. We also identify occasional transmission of daptomycin resistant organisms. Thus, control of daptomycin resistance in *E. faecium* may require interventions that both slow the emergence of resistance within patients and slow its spread. This work sheds light on the complex population dynamics leading to antibiotic resistance in hospitals.

## Introduction

The evolution of antibiotic resistance in a population includes both the origin of the resistance phenotype and its spread. Preventing the origin of resistance is often approached with higher doses or combinations of antibiotics ^1–3^, whereas controlling the spread of resistance, both within and between hosts, generally requires reducing selection by reducing antibiotic use and enhancing infection prevention ^4–6^. Characterizing the role of these two distinct processes is critical in situations where both may be acting simultaneously. For the important hospital pathogen vancomycin resistant *Enterococcus faecium* (VR *E. faecium*), resistance to daptomycin has been described to emerge within individual patients ^7–11^. Additionally, several studies have shown the existence of daptomycin resistant *Enterococcus* among patients without previous exposure to this antibiotic, which has been used to argue for the transmission of drug resistance^10, 12–14^. However, the frequency with which daptomycin resistance evolves within a patient, the genetic basis for daptomycin resistance, and how often resistance transmits between patients is not well characterized in the setting where *E. faecium* is endemic and resistance is increasing.

Previous studies have looked at the evolution of daptomycin resistance in patient derived isolates ^7,10,15–23^. These studies have identified many candidate resistance genes (Table S1), but the studies often included a small number of patients, and the population context of the antibiotic resistance evolution was not clear. A notable exception is a study that included 250 isolates from a single institution intermittently gathered over a 20-year period and included a genetically diverse set of *E. faecium* with a range of daptomycin resistance levels. Unlike other studies that identify potential daptomycin resistance mutations in a diverse range of genes, Wang *et al*. only identified mutations in *clsA* among the four patient having within-patient evolution in daptomycin resistance^10^.

The hospital system being studied here saw a rise in daptomycin resistance among blood stream infections with VR *E. faecium* ^8,9^. The current study seeks to clarify the underlying dynamics that led to the increase in daptomycin resistance. A case-control genomic comparison is performed with isolates obtained during a period of documented hospital wide increase in resistance and a subsequent decline when daptomycin use was limited ^9^. The goal is to determine how often resistance emerged *de novo* and whether there is any evidence of transmission of drug resistance between patients in the hospital.

The study finds that antibiotic resistance evolved many times, that the genetic pathways to resistance are also many, and finds support for occasional transmission of resistance between patients. The results give a clearer picture of the complex genomic and epidemiologic dynamics that lead to hospital-wide resistance patterns and highlight the multifaceted approach to antibiotic resistance management required in this setting.

## Materials and Methods

### Ethics

The work presented was approved by the Internal Review Board at the University of Michigan (ID number HUM00102282).

### Identification of cases and controls

Blood culture isolates were inconsistently stored in the clinical microbiology laboratory in the period from 2013 to 2016. From this convenience sample, we identified all patients with at least one daptomycin resistant *E. faecium* blood stream isolate, here defined as having a daptomycin minimal inhibitory concentration (MIC) greater than 4μg/mL, as measured in the clinical laboratory during patient care. In this period, the clinical microbiology laboratory used either E-test or the Trek method to measure the daptomycin MIC ^9^. Twenty-four patients with daptomycin resistant *E. faecium* were identified, 15 of whom had daptomycin resistance of their initial isolate (hereafter referred to as the R group of patients), and 9 had an initial susceptible isolate follow by a resistant isolate (hereafter referred to as the SR group). Previous infections were determined by searching for prior infection in the last 100 days.

For each of the 24 case patients, a control patient with *E. faecium* blood stream infection was identified from patients that never had a daptomycin resistant isolate reported and had an *E. faecium* isolate with a collection date that was the closest in time to the first blood stream isolate from each R or SR patient.

### Extraction of data

All doses of daptomycin administered in our hospital and the movement of patients through the hospital were extracted from the electronic medical record system.

### Antibiotic susceptibility testing

The MIC data reported by the clinical microbiology lab were extracted from the electronic medical record system. Susceptibility testing was repeated in the research laboratory using a broth micro-dilution method in calcium supplemented cation adjusted Muller Hinton broth using two-fold dilutions of daptomycin. Optical density at 600nm (OD) values were fitted to Hill function and resistance was quantified as the point at which the OD was inferred to drop to an OD indistinguishable from the blank as previously described ^11^.

### Sequencing

Whole genomic DNA preparations were submitted to the University of Michigan sequencing core for Illumina library preparation and paired end 125 bp Illumina Hi Seq2500, with 96 genomes multiplexed on a single flow cell (including isolates of *Enterobacter sp.* and *Pseudomonas aeruginosa*). For a subset of isolates, long read sequencing was also obtained using the MinIon (MIN-101B, Oxford Nanopore). Long read sequencing libraries were prepared using the SQK-LSK109 kit according to manufacturer protocols. Samples were multiplexed in two libraries and barcoded with the EXP-NBD104 Kit. The first library of 4 samples was run for 6 hours using a MIN-101Bsequencer on a MinION R9.4.1 flow cell. The flow cell was then washed using the EXP-WSH003 flow cell wash kit. The second library of 14 samples was then loaded and run for 24 hours. All 18 samples utilized unique barcodes.

### De novo assembly and annotation of genomes

Hybrid assembly using short and long read sequencing data was performed using Unicycler v0.4.4 using normal bridge setting ^24^. Multilocus sequencing types (MLST) were determined using MLST software against the pubMLST database (https://github.com/tseemann/mlst). Prokka was used to annotate the genome assemblies^25^.

### Phylogenetic reconstruction

A whole genome alignment was generated using Snippy (version 4.6.0), by aligning sequences to the strain DO chromosome^26^, then using gubbins (version 2.4.1)^27^ and RAxML^28^ to identify recombination and reconstruct the phylogenetic tree.

### Variant identification in candidate genes

A genome wide association study for variants contributing to daptomycin resistance would be dramatically underpowered with 82 clones because the number of genetic differences between isolates may be on the order of hundreds across the whole genome. We therefore focused on the candidate loci previously identify by Diaz *et al*.^17^, but excluded the hypothetical genes HMPREF0351_11658, HMPREF0351_10875, HMPREF0351_10830, HMPREF0351_12015, HMPREF0351_10297, HMPREF0351_10318, bacteriophage minor protein (HMPREF0351_10870), due to variable presence, lack of supporting data, or lack of plausible functional role. We included 50bp upstream or downstream of the candidate loci. These 36 candidate genes (table S1) comprise 44,837 bp, which is 1.66% of the strain DO chromosome. Short-read sequences from each isolate were mapped to the published sequence of strain DO (CP003583) ^26^ using the Burrows-Wheeler Aligner^21^. Variants were identified using the GATK variant calling pipeline ^29^. Identified variants were annotated with snpEff ^30^. Analyzed variants were further limited to regions in which there was sequence coverage (sequence depth >50) and the variant was present in at least 50 percent of the reads. All variants were individually visualize comparing the pileup of strains having and not having the variant using the IGV^31,32^. A pipeline that combines each of these steps in Snakemake ^33^ is available https://github.com/woodslab/between_hosts with dependencies and versions documented.

### Variant identification across the whole genome

Among closely related isolates, mutations were identified by mapping Illumina reads from subsequent strains to the *de novo* assembly of earliest isolates using the same workflow as above but using the *de novo* assembled genome as a reference, and utilizing the annotation generate by PROKKA ^25^. Structural variants were identified in comparisons where both strains had long-read sequence data using sniffles (v 1.0.11) ^34^.

### Data Availability

Sequences were deposited under NCBI Bioproject PRJNA746456.

### Visualization

Processing of output was performed using the Biopython libraries ^35^ and dendropy 3.5.3^36^. Phylogenetic trees were visualized using the ETE Toolkit 3.0.0b34 ^37^ and ggtree ^38,39^. Visualization of alignments was performed with Easyfig^40^.

## Results and Discussion

### Patient information

Within the available collection of isolates, 24 patients were identified with at least one daptomycin resistant *E. faecium* clinical isolate. Nine of these patients had an initial culture with a daptomycin susceptible isolate (here defined as MIC <= 4μg/mL), and then later a resistant isolate (here defined as MIC > 4 μg/mL). These patients were labeled SR1-SR9. The remaining 15 patients had daptomycin resistance of their initial isolate and were labeled R1-R15. Isolates from control patients, who never had a daptomycin resistant *E. faecium* blood stream isolate, were labeled C1-C24. In total, this study includes 48 patients with *E. faecium* bacteremia, including 24 of the 61 patients with daptomycin resistant *E. faecium* from June 2013 through January 2016.

### Incongruent measure of daptomycin resistance

The challenges in measuring daptomycin MIC have been extensively discussed previously, and may be due to differences in assay, genetic variability in the inoculum effect, sensitivity to calcium, or whether grown in biofilm ^9,41–43^. The MIC measured by repeated susceptibility testing using a micro-dilution method are shown (Figure 1). The relationship between clinical result, and the repeated testing using comparable 2-fold dilution are also shown (Figure S1), while they are correlated, the broth microdilution re-testing is typically lower in this study. In seven of the nine SR patients, the increase in resistance from the initial to the subsequent isolates was reproducible when remeasured (the two exceptions being SR3 and SR5). Additionally, six of the 15 R patients had isolates with laboratory measured susceptibility consistently above 4 μg/mL on retesting (Figure 1). Moving forward, we retain the R, SR and C labels, as this reflects the initial study design, which was determined by clinical designation of resistance and sensitivity. However, for the quantitative comparison of R and SR groups we will also report the results for the subsets of isolates that have resistance confirmed.

**Figure 1.**
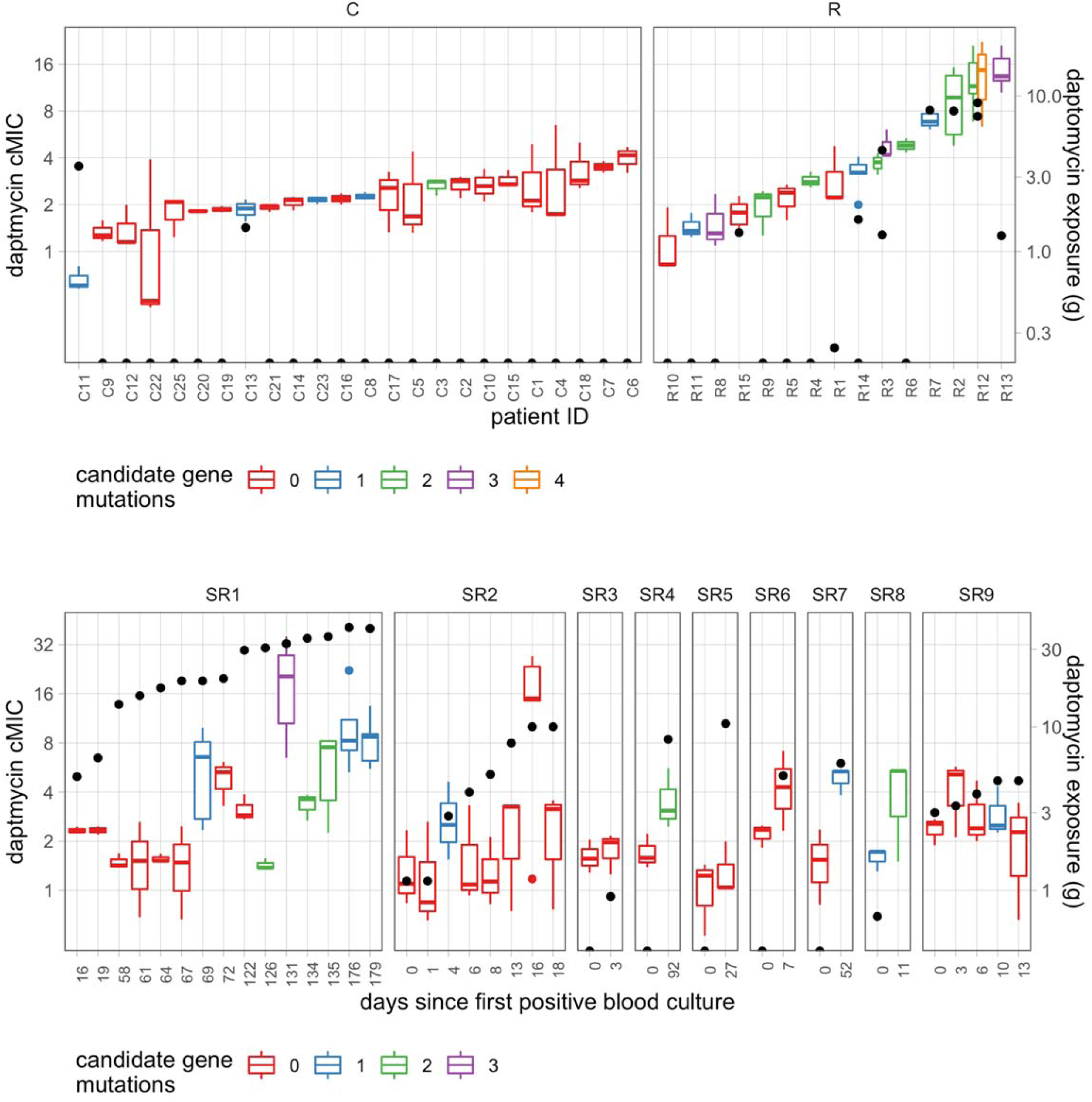
Resistance and candidate gene mutations in patients. The broth microdilution computed MIC (cMIC) for each of the Control patients (C) resistant patients (R) and the 9 SR Patients. Two of the patients, R3 and R12, had two isolates for sequencing, with different daptomycin exposure. The number of candidate mutations is indicated by color, and the amount of daptomycin administered in the prior 100 days is indicated by black dot.

### Phylogenetic clustering of R and SR patient samples

The maximum likelihood phylogenetic tree with recombination removed is presented (Figure 2). The sequenced isolates are a diverse sampling of the *E. faecium* clade A, including eight MLST types. All MLST types with more than one sequence contain both susceptible and resistant isolates. We can place a lower bound on the number of times daptomycin resistance arose independently by determining the fewest transitions from sensitive to resistant along this tree. This results in a minimum of 13 transitions among the 24 cases classified as resistant using the clinical laboratory designation of MIC > 4μg/mL. Among the 13 patients with resistance confirmed in the subsequent broth microdilution assay, there were at least 10 independent evolutionary events. In 8 of the 9 SR pairs, the strains are very closely related, consistent with with-host evolution. The remaining pair, from patient SR4, the two sequences were from separate MLST types, indicated two independent introductions into this patient.

**Figure 2.**
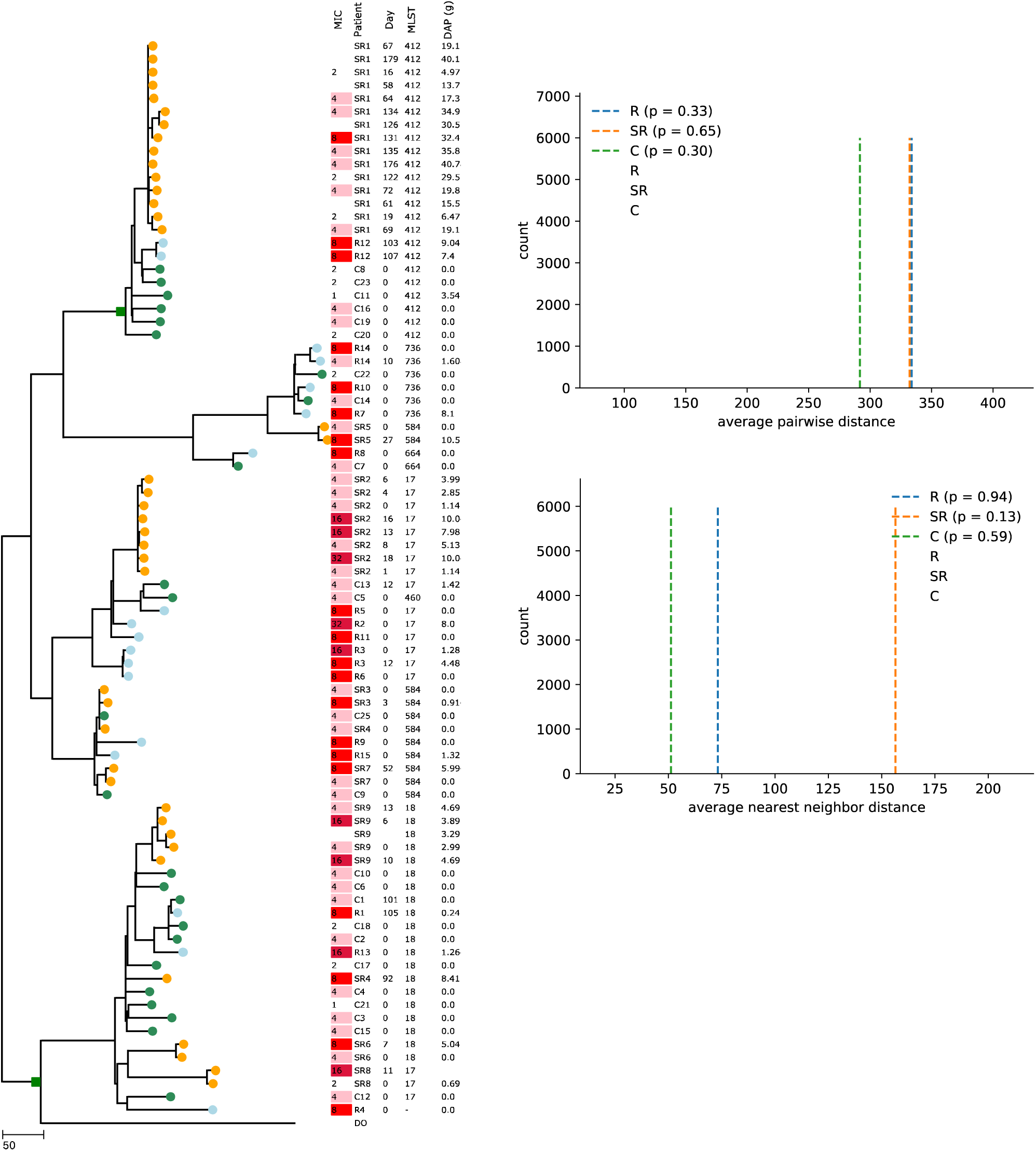
Core genome phylogeny. Core genome phylogeny with recombination removed of 82 isolates in this study. Isolates from SR patients are in orange, R patients in pink and C patients in blue. Scale bar indicated SNP distance. ‘Day’ indicates the days from the first isolate, ‘MIC’ is the MIC identified in the clinical microbiological lab, and the daptomycin exposure in the 100 days prior to isolate collection (in grams) is indicated. Gubbins analysis identified two putative recombination events involving the LiaFSR locus: two gains of the Clade B like allele indicated with green squares. The results of the phylogenetic clustering analysis is indicated in panels B and C, where the point estimate is indicated with a vertical dashed line, and the distribution base on bootstrap randomization is indicated by the histogram. The null is not rejected for any of the comparisons, thus there this analysis does not identify phylogenetic clustering.

Phylogenetic clustering of the R and SR patients was assessed in two ways, using the phylogenetic net relatedness index (NRI) and nearest taxa index (NTI) of Webb ^44^. To calculate these indices, the phylogenetic tree (Fig 2) was further trimmed to only include the first isolates per patient and remove patient SR4, as the two isolates of this patient were phylogenetically distinct (repeating the analysis with SR4 kept in the analysis had no impact on the significance). Comparison of the observed NRI and NTI values to 100,000 bootstrap values, obtained by randomly reassignments of the tips of the tree, show that they did not differ from this random expectation (Figure 2). Thus, this analysis yields no evidence for phylogenetic clustering of the resistant isolates.

### Mutations in the candidate loci

We next turned to identifying the genetic changes that account for the daptomycin resistance phenotype by looking at variants in candidate daptomycin resistance genes across all sequenced isolates. The variant calling approach identified 352 mutations and 93 unique protein altering variants in 24 candidate genes when compared to reference strain DO^17^ (Table S2). Most variants were synonymous, consistent with these candidate genes being conserved over most of the evolutionary history represented across the sample collection. Most of the specific mutations in candidate genes have not been previously characterized. Some mutations were shared among multiple MLST types in the sample, suggesting either a more distant origin or convergent evolution at the genetic level. Because the phenotypic assessment showed that the resistance was emerging within MLST types, and no MLST types showed a predisposition for resistance (*i.e.* there was no phylogenetic clustering of resistance), we removed all the variants that were shared across all isolates from multiple MLST types, to focus on mutation that accounted for the conversion to daptomycin resistance.

Overall, 11 candidate genes contained variants that met our criteria for mutations in candidate loci using the above definition (Table S3), including 6 different candidate genes being mutated between the sensitive and resistant isolates of SR patients (Table 1).

**Table 1.**
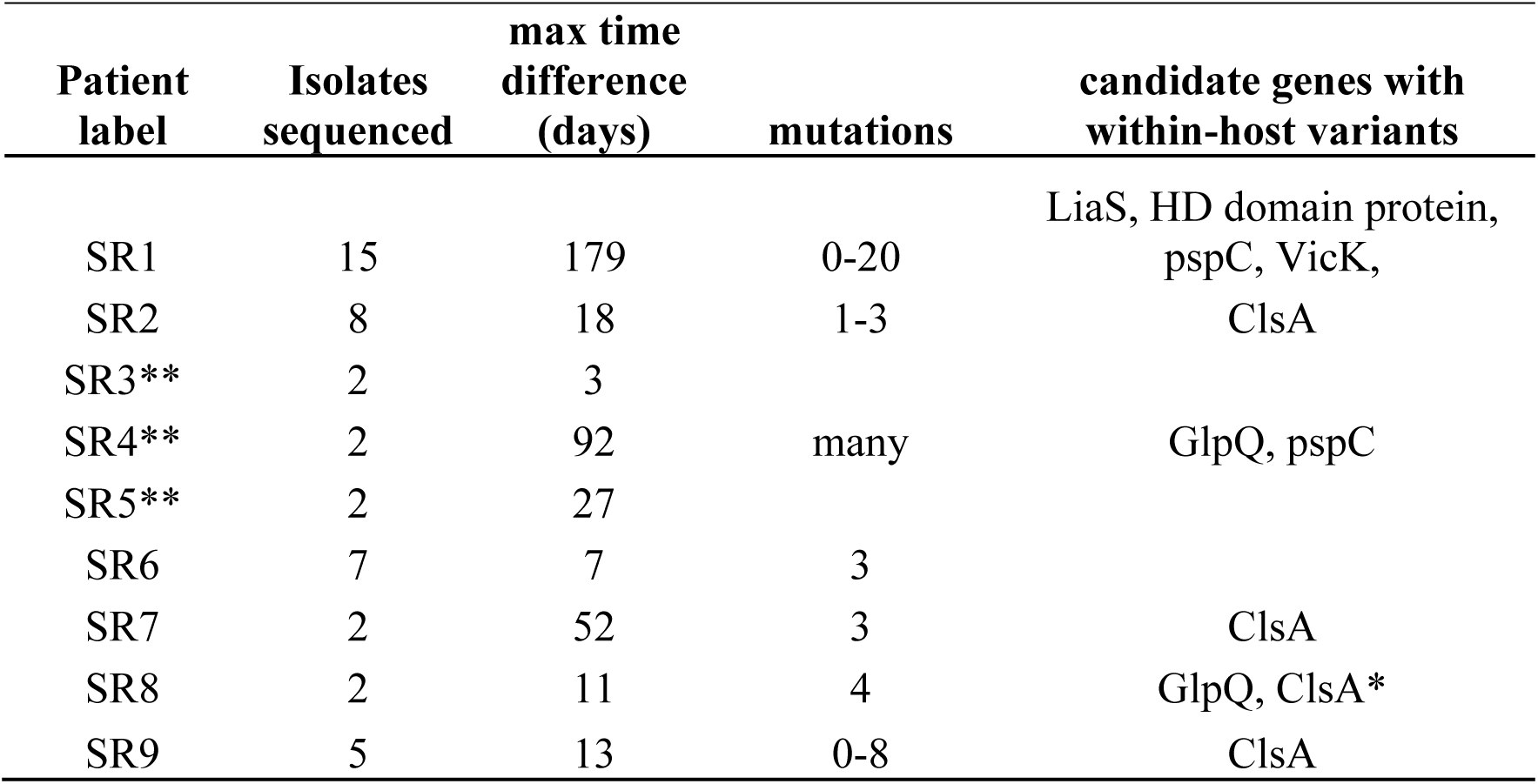
Summary of within host variants in SR patient. * The ClsA mutation (p.Gly408Ala) was present in the sensitive isolate from patient SR8 as well as the resistant isolate. ** Whole genome comparisons were not performed for patient SR3 and SR5 because there was no reproducible change in daptomycin susceptibility between isolates, or in patient SR4 because isolates were of different MLST types.

### Mutations in the cardiolipin synthase A

The *clsA* gene is the candidate gene most frequently containing mutations (Table S3), multiple potential mutations were observed, and mutations in *clsA* do not by themselves necessarily result in an MIC > 4. Four of the nine SR patient have isolates with mutation in ClsA, D27N in SR2, R218Q in SR7, H215R in SR9, and both isolates of SR8 have the mutation G398A. Additionally, six of the 15 patients in the R group had ClsA mutations: C267R in R3a, L22 in R8, A20P in R8, H141Y in R13, V37A in R2, N13I in R11, V37A in R12. Only one control patient, C11, has a mutation in this gene, which was D27N.

### Mutations in the *liaFSR* operon

Among all isolates in this study, the *liaFSR* operon sequence was either very closely related to the reference strain DO, or they differed at 23 amino acid: LiaF:R7K, H55Y, A86V, F88L, I119V, V156I, Y224C, I243V, LiaS:K4R, R7K, L57I, T77A, L98F, L100H, L101H, K117M, T120A, V136I, S258T, and LiaR:T45A, W73C, E75K, E142D. These two divergent alleles shared 93% nucleotide identity and 97% amino acid identity across the three genes. This level of difference is similar to the divergence identified between clade A and clade B of *E. faecium*^45^. The divergent alleles were compared to a clade B strain com12 (ASM15763v1, genes EFVG_02032, EFVG_02032,EFVG_02031) ^45^ which revealed strain DO was 99% identical to com12 at the nucleotide sequence and 100% for the amino acid sequence (Figure 3). The Com12-like allele of strain DO is shared in all ST412, ST18, and some ST17 in our dataset, consistent with horizontal gene transfer between clade A and clade B at some point in the past. The gubbins analysis identified two putative recombination events involving the *liaFSR* operon, consistent with two gains of com12 like allele (green squares in Fig. 2). Given the challenge of constructing a core genome phylogeny in the face of frequent recombination ^46^, the number and phylogenetic position of these horizontal gene transfer events should be viewed as hypotheses requiring subsequent validation. Visual inspection with the alignment of this region between Com12 and DO, suggest a horizontal gene transfer event containing the entire *liaFSR* operon (Figure 3).

**Figure 3.**
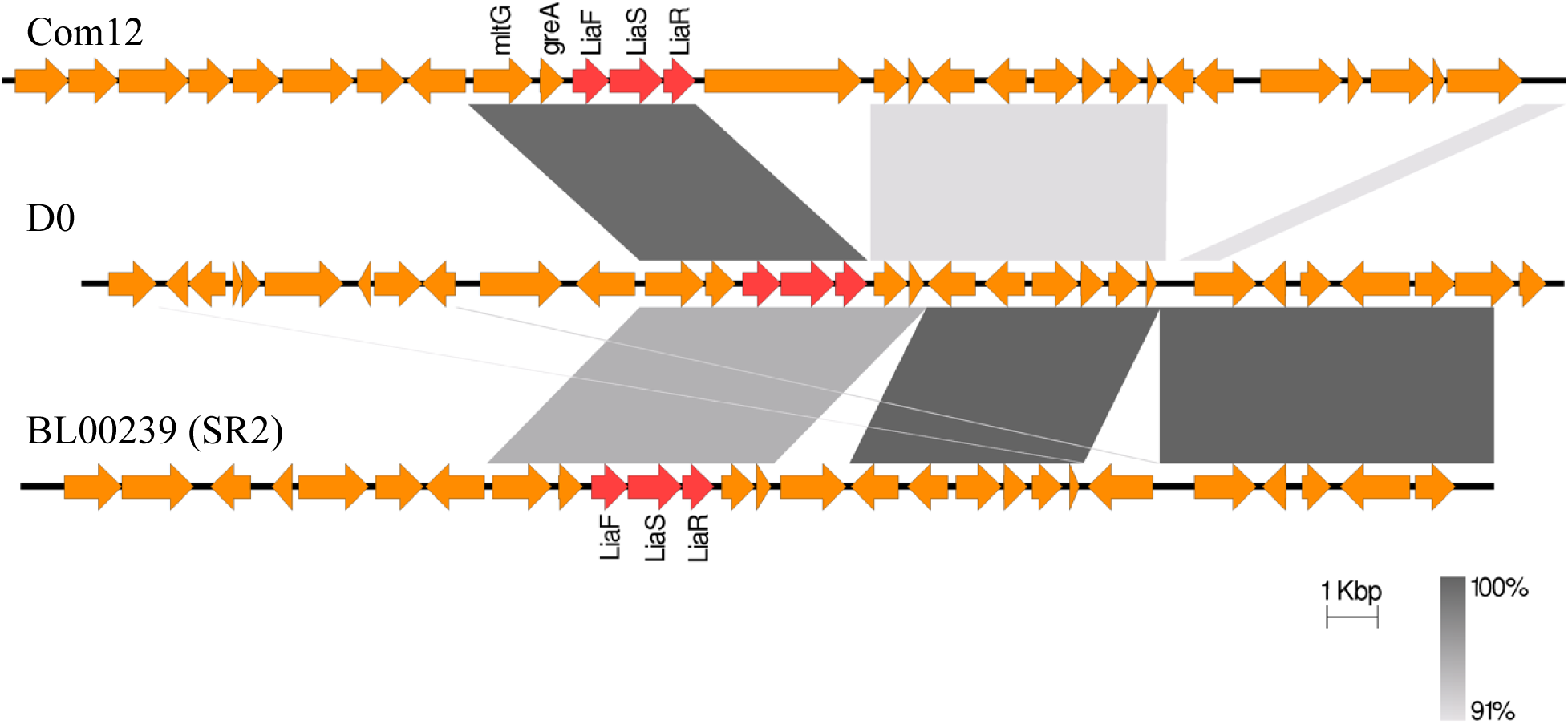
LiaFSR alignment. Alignment of portions of the genome of isolates Com12 (top), DO (middle), and the initial sensitive isolate from patient SR2 (bottom), which show that the LiaFSR locus of strain DO is nearly identical to the Clade B isolate Com12, and differs from the other Clade A isolates (here represented by the sensitive isolate from patient SR2) by about 7%. This occurs in a region with limited homology between DO and other isolates.

Previous studies have focused on the mutations in LiaR (W73C) and LiaS (T120A)^17^ and were cited among the reasons for changing clinical break points for daptomycin for *E. faecium* ^47^. These two mutations are among the 23 included in putative horizontal gene transfer event between Clade A and Clade B of *E. faecium*, a divergence previously estimated to have occurred over 2000 years ago ^48^. Thus, the previously hypothesized co-evolution ^17,49^ between these two codons may have occurred once, over a long period of time and in the context of multiple changes at other sites. There is an association between the LiaR73C and LiaS120A containing allele with a higher MIC, and concordantly the number crossing the threshold of daptomycin resistance (Fig. S2). However, linkage with many other variants within this operon and elsewhere in the genome preclude attributing phenotypic changes to these two specific amino acid changes using the comparative approach of the present study.

In addition to the difference between these two divergent alleles, four additional mutations were seen in this operon, but each was seen in one patient only and all occurred on the Com12-like background. LiaF V190L was identified in R13, LiaS A88V in R4, and LiaS V129G, which arose in patient SR1. In a later isolate from this same patient, SR1, a LiaS E192* nonsense mutation in addition to the LiaS V129G mutation. The LiaF V190L, LiaS A88V, and LiaS V129G, mutations were seen in isolates with elevated MIC, (16, 8, and 8 respectively), but the strain with the combined LiaS V129G E192* has an MIC of 4.

Previously identified mutations in the LiaR cell wall stress response regulator protein have been associated with increase in activity of LiaR and increased daptomycin resistant, whereas deleting LiaR results in increased daptomycin susceptibility ^50^. Two amino acid changes have been shown to increase the activity of the gene product ^51^. In one study, *in vitro* selection in the presence of daptomycin results in a LiaS G92D mutation, but the subsequence propagation in the absence of antibiotic results in insertion sequence mediate disruption of the response ^52^. The evolution in patient SR1, described here, is consistent with that experimentally observed pattern, with a potentially activating mutation leading to resistance and a followed by a reversion with a mutation that may abolishes the regulatory function of the operon all together. However, this *in vivo* reversion is more complex, as it also does not carry the mutations in the HD domain protein nor walK, indicating that observation of a lower resistance in the later isolate may be multifactorial.

### Whole genome analysis of within host evolution in SR patients

In the six patients in whom the increased daptomycin resistance was detected by clinical testing and was also confirmed using repeated MIC testing by broth microdilution, we obtained long read sequencing to close or nearly close the genome. Patient SR4 was not included because the two isolates were of different MLST types. Long read sequencing was obtained for the earliest isolate, and the most resistant isolates. A total of 91 unique variants were identified within hosts. Of these, 77 were in coding regions, 10 of these were synonymous while the remainder were non-synonymous (50), frameshift (10), nonsense (4) or otherwise changed the expressed coding sequence (3) (Tables S4-S15). Most of these changes are of unclear significance, but some mutations and patterns warrant comments.

First, isolates from patient SR9 have a mutation in RecX. RecX interacts with the recombinase RecA and may result in decreased activity (in *E. coli*). This is notable because, RecA has been suggested to be important to physiological responses to antibiotics ^53^. In the *in vitro* setting, strains with RecA mutation evolved linezolid resistance more slowly, the mechanism was suggested to be slower gene conversion ^54^, and has been suggested to be a target of adjuvant chemotherapy to slow the emergence of resistance^55^. Notably, mutations in recombination pathways have been described in daptomycin resistant *E. faecium* previously^19^.

Second, a common component of within patient evolution, is non-synonymous changes in membrane transporters. Whether this reflects changes in the nutrient environment in the transition to blood stream, response to antibiotics, potential immune selection ^56^ or other microevolutionary pressures, such as bacteriophage or bacteriocins that reside in the complex biotic and abiotic environment is unclear at this point.

### Parallel mutations across SR patients

Apart from the candidate genes, there were four genes mutated in more than one SR pair. Resistant isolates from patient SR2 and SR6 gain mutations in or near the CroRS two-component signal transduction system. CroS sensor kinase and its cognate response regulator CroR (HMPREF0351_12687 and HMPREF0351_12688 on strain DO respectively^57^) have been shown to be responsible for cell wall stress response and critical for beta-lactam resistance in both *E. faecalis* ^58,59^ and in *E. faecium* ^57^. The SR2 resistant isolate has a complex inversion, with a break point 743 bases upstream of the CroR start. Patient SR6 contains the variant N100T in CroR.

Isolates from patient SR6 and SR2 also have disruptive mutations in a gene annotated as *yhhT* by Prokka, which encodes an AI-2E family transporter. SR6 contains a transposase insertion into this gene, and one isolate from SR2 (BL00221-1) gains a stop codon at position 115. Disruptive mutations in this AI-2E family transporter were associated with daptomycin resistance obtained by in vitro selection in *Staphylococcus aureus* (locus tag SAUSA300_RS04910)^60^.

The folate transporter FolT is mutated in two patients, FolT I33II in patient SR6, and FolT T121A in patient SR8. The T121A mutation was studied in the *E. faecalis* sequence, and found to decrease binding efficiency by 10 fold^61^ which may be significant, as most *E. faecium* are thought to require folate ^62^. The I31 codon deletion is in the L1 loop region ^61^.

The hypothetical gene corresponding to HMPREF0351_11546 in DO was mutated in patient SR1 and patient SR9, both are frameshift mutations. Whether this or any of these specific parallel mutations contribute to daptomycin resistance would require additional investigation.

### Evidence for transmitted resistance

Transmitted resistance is suggested when a patient has a resistant isolate as their first isolates but had no previous exposure to daptomycin. This was the case in 8 of the 15 R patients (R4, R5, R6, R8, R9, R10, R11 and R14), although only one of these patients (R6) consistently had a resistance above 4μg/mL upon re-testing (Figure 1). Additionally, all isolates with an MIC of 16 or 32 had previous daptomycin exposure raising the question of whether higher level resistant strains have reduced ability to either transmit or establish subsequent infections.

In two pairs of patients there is stronger evidence that transmission of resistance has occurred, consisting of phylogenetic and epidemiological links (Figure 4). The isolate from patient R8, is most closely related to the isolate from patient C7, differing by 15 mutations, suggesting that the two candidate mutations (ClsA A20P, and GlpQ V595L) could have arisen in patient C7 during treatment, or during a recent transmission chain within the unit (Figure 4). Second, patient R6 is closely related to isolates from patient R3, and two candidate mutation are shared between these isolates, strongly suggesting that this represents transmission of a resistant isolate between patients.

**Figure 4.**
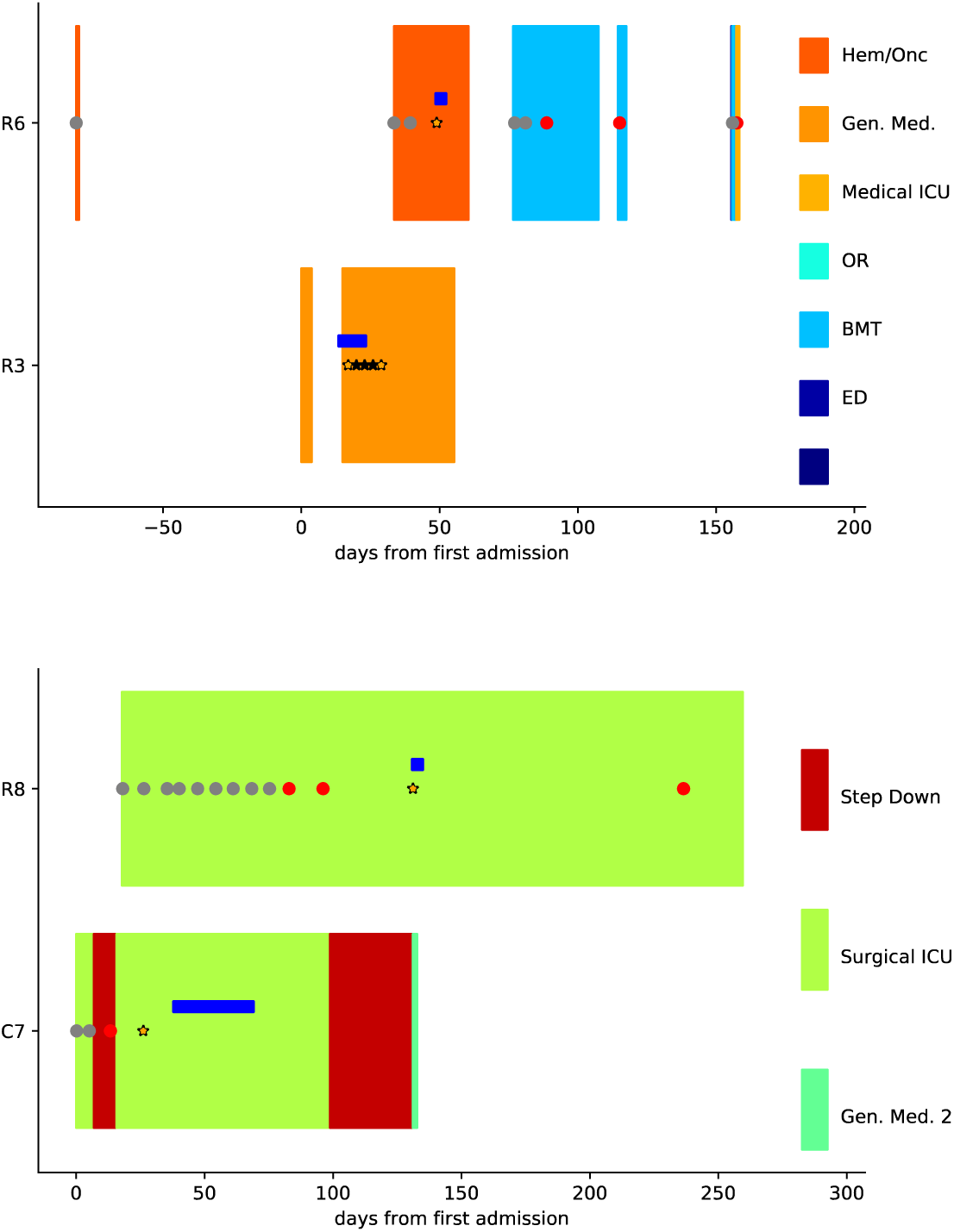
Putative cases of transmitted daptomycin resistance. Two possible transmission clusters show patients overlap in the hospital unit, at the same time, in only a subset of closely related strains. Each hospital unit is given a unique color. VRE surveillance culture using peri-rectal swabs are indicated in circles which are red if positive and grey if negative. Positive blood cultures are indicated with stars, from which sequencing were obtained. Dark blue horizontal bars indicate daptomycin treatment.

It is often proposed that antibiotic use in one patient may lead to resistant infection in another patient, although concrete examples are vanishingly rare. The sequencing data for patient C7 and R8 are most consistent with this scenario: C7 and R8 are epidemiologically linked giving opportunity for transmission, plus the most-recent common ancestor could plausibly be in C7 as the sequence of this isolate was only one mutation from the most recent common ancestor, using the DO sequence as the outgroup. Patient R8 had no prior daptomycin exposure, but patient C7 was exposed to daptomycin before the two mutations associated with daptomycin resistance were seen in R8.

## Conclusion

Taken together these result show that the evolution of daptomycin resistance was remarkably dynamic in this hospital-associated population of *E. faecium*. Resistance was gained and lost within patients, across a range of genetic backgrounds, and utilized multiple different genetic pathways.

The genetic diversity across the studied population was high. Eight different MLST types were identified, with at least one resistant isolate in each of these MLST types, and seven also have susceptible isolates. Thus, the trend of increasing daptomycin resistance observed at the hospital level from 2013-2016 ^8,9^ was not due to the cryptic spread of a single daptomycin resistant clone, but instead was driven by frequent *de novo* evolution and occasional transmission between patients. There is evidence for transmission of resistant strains between patients based on the 8 patients with resistance on their first isolate without prior use of daptomycin use in our records, and two patient-pairs with epidemiological and phylogenetic evidence for transmitted resistance, R3-R6 and C7-R8.

Numerous studies have previously looked at the evolution of daptomycin resistance in patient derived isolates^7,10,15–20^ and have shown that an array of mutations can be associated with daptomycin resistance. These mutations were summarized by Diaz *et al.* ^17^. Most of these candidates have not been experimentally validated apart from the major cardiolipin synthase, and some mutations in the LiaFSR operon. The current study further supports the role of these candidates as targets of selection under daptomycin pressure, with emphasis on the major cardiolipin synthase, LiaFSR operon, an HD domain protein, and the yycHIJ operon. The results of this study validate the work of others that clinically observed within-host evolution is compatible with *de novo* emergence of resistance while being treated with daptomycin. However, this study differs from Wang *et al.* ^10^, which only identified ClsA mutations among its 4 SR pairs.

Limitations of the current study include that only a single center is represented and that sampling was incomplete. It is likely that the antibiotic exposure and the frequency of transmission will vary between centers based on community prevalence, infection prevention practices, and antibiotic utilization practices^63^. However, this is the first study to describe the genetics of transmission and resistance evolution in the context of well-described recent population-wide increase in resistance^8,9^. Additionally, this study suffers from the challenge of quantifying daptomycin resistance. Many of the isolates that were identified as resistant based upon susceptibility testing in the clinical laboratory were not confirmed to be resistant with the assays used in the research laboratory. While the MIC discrepancy impacts the quantitative results, the same general conclusions hold regardless of the resistance assay used. Finally, we describe two novel candidate daptomycin resistance targets (CroSR and AI-2E family transporter in *E. faecium*) which deserve additional investigations, but should be considered preliminary as selection for reasons other than daptomycin or non-adaptive process such as drift or hitchhiking may be at play.

How to use antibiotics best to slow the emergence of resistance remains controversial ^64^. Recent theoretical work and results from model systems have suggested that preventing *de novo* acquisition of resistance may be best achieved with aggressive antibiotic treatment aimed at preventing mutations from occurring in the first place, whereas reducing selection on preexisting resistance variants may be better accomplished with minimal treatment, which limits the selective pressure on the spread of existing resistant variants ^4–6^. For daptomycin, new guidelines suggest that higher doses may be used ^47^, with a goal of preventing the emergence of drug resistance over the course of a patient’s infection. These recommendations are in part based on *in vitro* and animal model data that suggest these doses can prevent the emergence of resistance^65,66^. However, the results presented here suggest the need to balance the goal of preventing resistance at the site of infection (e.g. blood stream), with the goal of reducing the risk of resistance emergence and onward transmission, which presumably arises in the colonizing population in the intestinal tract ^11^. Optimal solutions for one are not necessarily optimal solutions for the other. Novel approaches to isolate the selective forces in each compartment are promising^67^. The results presented here point to the need to identify approaches that balance treatment of transmitted and *de novo* evolved resistance.

## Supplemental data

**Table S1** – Table of candidate genes associated with daptomycin resistance.

**Table S2** - List of all mutation in candidate genes relative to reference strain DO across the 82 sequenced isolates.

**Table S3** – List of candidate gene mutations the were after removing those that do no alter protein coding and regions or those that do not (Candidate_gene_mutations_filtered.doc)

**Table S4** – SR1 within host sort_read_mutations_believed

**Table S5** – SR1_structural_variants_believed

**Table S6** – SR2_sort_read_mutations_believed

**Table S7** – SR2_structural_variants_believed

**Table S8** – SR6_short_read_mutations_believed

**Table S9** – SR6_structural_variants_believed

**Table S10** – SR7_short_read_mutations_believed

**Table S11** – SR7_structural_variants_believed

**Table S12** – SR8_short_read_mutations_believed

**Table S13** – SR8_structural_variants_believed

**Table S14** – SR9_short_read_mutations_believed

**Table S15** – SR9_structural_variants_believed.docx

---

**Figure S1.**
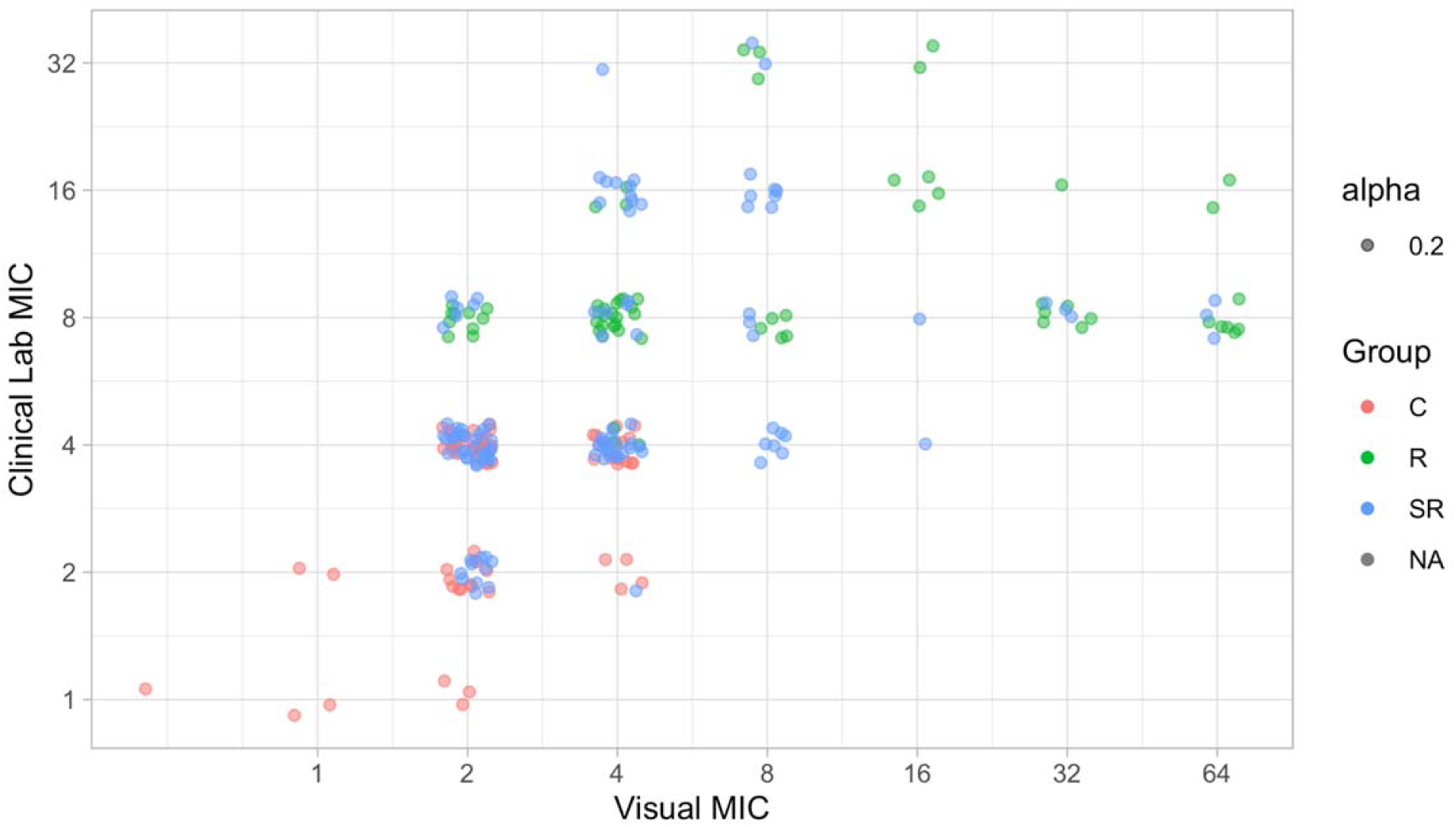
The visual MIC on 3 independent replicates for each of the 82 isolates. Both visual MIC and the Clinical Lab MIC were measures in discrete 2-fold values the plot adds artificial ‘jitter’ to better visualize samples at the same values.

**Figure S2.**
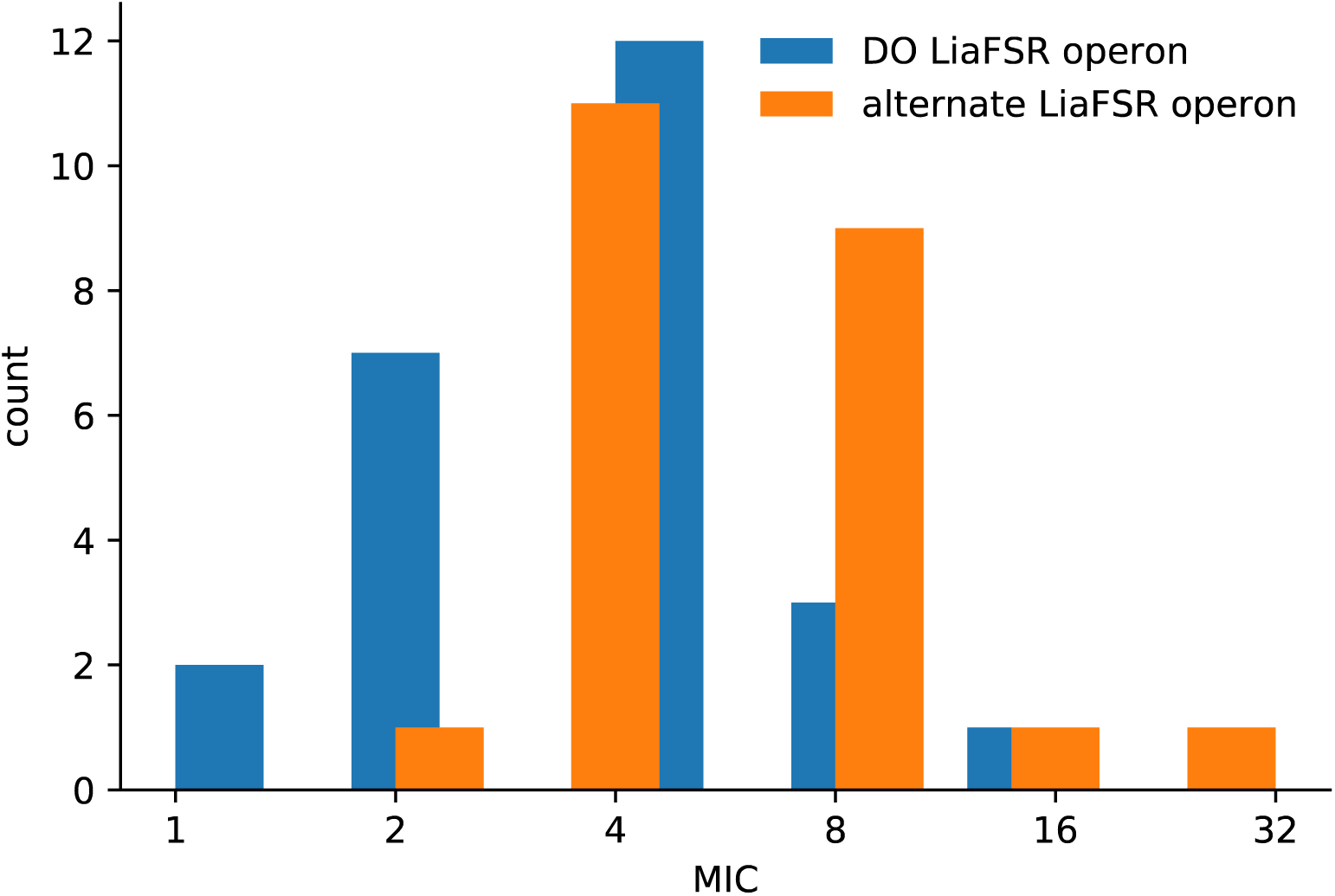
The distribution of daptomycin MIC values among the isolates containing the DO-like allele of the LiaFSR operon containing the LiaR73C and LiaS120A amino acid variants, or the alternate allele.

